# LOSS OF ROR2 TYROSINE KINASE RECEPTOR IS ASSOCIATED WITH ENDOTHELIAL DYSFUNCTION IN PAH VIA INAPPROPRIATE INTEGRIN β1 ACTIVATION

**DOI:** 10.1101/2025.04.28.651135

**Authors:** Ankita Mitra, Ananya Chakraborty, Brian Zhong, Lyong Heo, Stuti Agarwal, Amanda Pacheco, Natasha Auer, Alexander Dunn, Prakash Chelladurai, Ananya Jain, Juan Andrés Muñoz Matos, Anuradha Bankar, Eleana Stephanie Guardado, Dan Yi, Hanqiu Zhao, Kwun Wai Dede Man, Ramesh Nair, Olivier T. Guenat, Zhiyu Dai, Vinicio A. de Jesus Perez

**Affiliations:** Division of Pulmonary and Critical Care, Stanford University, Palo Alto, CA, United States; Department of Chemical Engineering, Stanford University, United States; Stanford Center for Genomics and Personalized Medicine, Stanford University School of Medicine, United States; Department of Internal Medicine, University of Arizona College of Medicine Phoenix, AZ; Division of Pulmonary and Critical Care and Medicine, Department of Medicine, School of Medicine, Washington University in St. Louis, St. Louis, MO.; ARTOG Center for Biomedical Engineering Research, University of Bern, Switzerland

**Keywords:** Pulmonary Hypertension, Angiogenesis, Wnt signaling, Endothelium, Focal Adhesion, Integrin, Vascular Biology

## Abstract

**Rationale:** Endothelial dysfunction is a key feature of pulmonary arterial hypertension (PAH). We previously identified Wnt7a, a ligand of the Wnt planar cell polarity (PCP) pathway, as essential for pulmonary angiogenesis, with its loss linked to PAH. Given the importance of Wnt/PCP to lung endothelial function and angiogenesis, our goal is to elucidate how Wnt/PCP regulates angiogenic responses in pulmonary microvascular endothelial cells (PMVECs). ROR2, a tyrosine kinase receptor specific to Wnt/PCP, is crucial for cardiovascular development, but its role in PAH is unclear. We hypothesized that ROR2 supports endothelial homeostasis, and its loss would impair angiogenesis, contributing to PAH.

**Methods:** Endothelial-specific ROR2 knockout (ROR2 ECKO) and wild-type (WT) mice were studied under normoxia and chronic hypoxia using echocardiography, hemodynamics, and lung morphometry. PMVECs from healthy and PAH lungs were transfected with ROR2 siRNA/constructs for functional and molecular studies. Focal adhesion (FA) activation and force generation were assessed via FRET-based methods. Bulk and single-cell transcriptomic analyses were performed on siROR2 PMVECs and ROR2 ECKO lungs.

**Results:** ROR2 ECKO mice exhibited worsened pulmonary hypertension, right ventricular remodeling, microvascular loss, and muscularization in hypoxia. Single-cell RNA sequencing of lung endothelial cells showed dysregulation of pathways involved in barrier formation and angiogenesis. Evans blue dye extravasation confirmed reduced endothelial barrier integrity in ROR2 ECKO mice. ROR2-deficient PAH PMVECs displayed increased adhesion, permeability, and FA numbers, with reduced VE-cadherin at cell junctions. Confocal imaging revealed ROR2 localization in FAs, interacting with integrin β1 (ITGB1). FRET analysis showed that ITGB1 remained in an active, adhesion-promoting state in ROR2-deficient cells. Restoring ROR2 in PAH PMVECs normalized adhesion, barrier function, and FA abundance. Transcriptomic analysis identified Rab12 as a key mediator of ROR2-ITGB1 crosstalk, with Rab12 knockdown mimicking ROR2 deficiency in PMVECs.

**Conclusions:** ROR2 regulates pulmonary angiogenesis by maintaining endothelial barrier integrity and facilitating integrin recycling. Restoring ROR2 signaling could be a potential therapeutic approach for PAH.

## INTRODUCTION

Pulmonary arterial hypertension (PAH) is a life-threatening disorder characterized by increased pulmonary pressure and right heart failure^1,2^. A hallmark of PAH is the alteration in the structure and function of lung capillaries, where vessels show increased leakiness, abnormal morphology, and remodeling^3^. Current PAH therapies fail to prevent disease progression due to the inability to halt the loss or promote the regeneration of lung microvessels^4,5^. A significant roadblock to developing novel therapeutics is an incomplete understanding of the mechanisms responsible for pulmonary endothelial dysfunction and impaired regeneration of damaged vessels. Understanding the underlying mechanism that regulates lung angiogenesis is imperative, as it will accelerate the discovery of new therapies.

In healthy individuals, endothelial cells regenerate lost or damaged vessels through angiogenesis, which occurs in two phases: 1) angiogenic sprouting and 2) endothelial barrier establishment^6,7^. Our group has shown that the Wnt/planar cell polarity (PCP) pathway, a developmental pathway responsible for coordinating cell motility and tissue morphogenesis, promotes angiogenic activity by pulmonary microvascular endothelial cells (PMVECs)^8–10^. Recently, we showed that the Wnt/PCP ligand Wnt7a regulates sprouting angiogenesis through its capacity to enhance VEGF signaling in PMVECs^11^. Using both *in vitro* and *in vivo* models, we demonstrated that Wnt7a is required for sprouting angiogenesis, and its loss in PAH contributes to endothelial dysfunction and vascular remodeling.

A central question arising from this work was the mechanism by which Wnt7a activates Wnt/PCP signaling in PMVECs. A genetic screen of Wnt receptors identified ROR2, a tyrosine kinase receptor specific to Wnt/PCP, as the primary candidate for serving as a Wnt7a receptor. ROR2 is involved in cardiovascular development^12,13^, and its deficiency has been linked to congenital cardiovascular disorders such as tetralogy of Fallot^14^. In our studies, we confirmed that the knockout of ROR2 reduces the biological response of healthy PMVECs to Wnt7a and prevent angiogenic sprouting. In PAH PMVECs and in vascular lesions, we confirmed that ROR2 expression was significantly reduced^11^. However, beyond these provocative data, little is known about the full scope of angiogenic activities of ROR2 in pulmonary endothelium and the mechanisms by which ROR2 deficiency can contribute to PAH.

In this study, we examined the contribution of ROR2 in PMVECs and how disruption of ROR2 expression affects lung angiogenesis beyond the sprouting stage. We hypothesized that ROR2 activation in pulmonary microvascular endothelial cells (PMVEC) is required for pulmonary angiogenesis, and its loss may be associated with PAH development. We found that ROR2 expression is significantly reduced in vascular lesions and PMVECs from PAH patients and that ROR2 knockdown reduces endothelial barrier integrity through an increase in adhesion and junctional disassembly. We defined a novel mechanism that positions ROR2 as a critical modulator of ITGB1 activation through the engagement of “inside-out” signaling processes that rely on the crosstalk between ROR2 and ITGB1. Thus, we show for the first time to our knowledge that ROR2 is a master regulator of lung angiogenesis and provides a novel paradigm to understand the pathogenesis of endothelial dysfunction and vascular remodeling in PAH.

## METHODS

An expanded materials and methods section containing a detailed description of reagents, techniques and assays is provided in the data supplement.

### Animals

*Mice:* All experimental protocols used in this study were approved by the Animal Care Committee of Stanford University and adhered to the published guidelines of the National Institutes of Health on the use of laboratory animals. C57BL/6J *ROR2^flox/flox^* mice were generated as previously described^15^. C57BL/6J VE-Cadherin (PAC)-CreERT2 mice were a kind gift from Dr. Ralf Adams (Max Planck University)^16,17^. Endothelial cell specific ROR2 knockout mice (hereafter referred to as ROR2 ECKO mice) were obtained by crossing *ROR2^flox/flox^* and VE-Cadherin (PAC)-CreERT2 mice for at least 5 generations. Ear clip-based genotyping of the progeny was used to identify littermate controls and ROR2 ECKO mice. Mice were housed under a 12-hour light and dark cycle with free access to food and water, under pathogen free conditions. Mice were treated with 20 mg/mL tamoxifen (dissolved in corn oil) for 5 consecutive days to induce the ROR2 knockout in all transgenic stains. After tamoxifen treatments, mice were allowed to rest for 7-10 days prior to experimentation.

### Hypoxia Studies

In experiments involving chronic exposure to hypoxia, mice were placed in a hypoxia chamber and exposed to 10% inspired O_2_ with access to food and water ad libitum for up to 3 weeks. The chamber was regulated with a continuous mixture of room air and nitrogen. The chamber environment was monitored using an oxygen analyzer (Servomex, Sugar Land, TX), and was inspected daily for O_2_ concentration, CO_2_ concentration and animal welfare. Chamber temperature was maintained at ambient room temperature (22-24°C). At the end of the treatment period, animals were euthanized using controlled isoflurane flow and cervical dislocation. The mice were then exsanguinated by perfusion through the right ventricle with 15-20 mL of 1X PBS at 37°C. Lungs and hearts were harvested and processed for OCT embedding.

### Statistical Analysis

The number of samples or animals studied per experiment is indicated in the figure legends. Values from multiple experiments are expressed as mean +/- SEM. Statistical significance was determined using unpaired t-tests or one-way ANOVA followed by Dunnett’s or multiple comparison tests unless stated otherwise. A value of p<0.05 was considered significant.

## RESULTS

### Loss of ROR2 is associated with severe pulmonary hypertension and vascular remodeling in mice

To test whether endothelial ROR2 insufficiency can contribute to pulmonary vascular remodeling *in vivo*, we generated a tamoxifen-inducible ROR2 ECKO mice and exposed them to normoxia vs. hypoxia (**Fig. 1A**). While there were no differences between wild type (WT) and ROR2 ECKO in normoxia, we observed significantly greater right ventricular systolic pressure (RVSP, **Fig. 1B**) and right ventricular remodeling (**Fig. 1C**) in ROR2 ECKO vs. WT mice under hypoxia. Echocardiographic assessment of the cardiac chambers revealed that hypoxic ROR2 ECKO demonstrated greater RV dilation vs. WT (**Fig. 1D**), consistent with the higher pulmonary pressures and RV remodeling seen in hypoxia (see **Fig. 1B-C**). Echocardiographic measurements of RV function revealed that RV fractional shortening was reduced in ROR2 ECKO mice in hypoxia **(Fig. 1E)** whereas the pulmonary artery velocity time integral (PA VTI) did not change compared to WT (**Fig. 1F**).

**Figure. 1:**
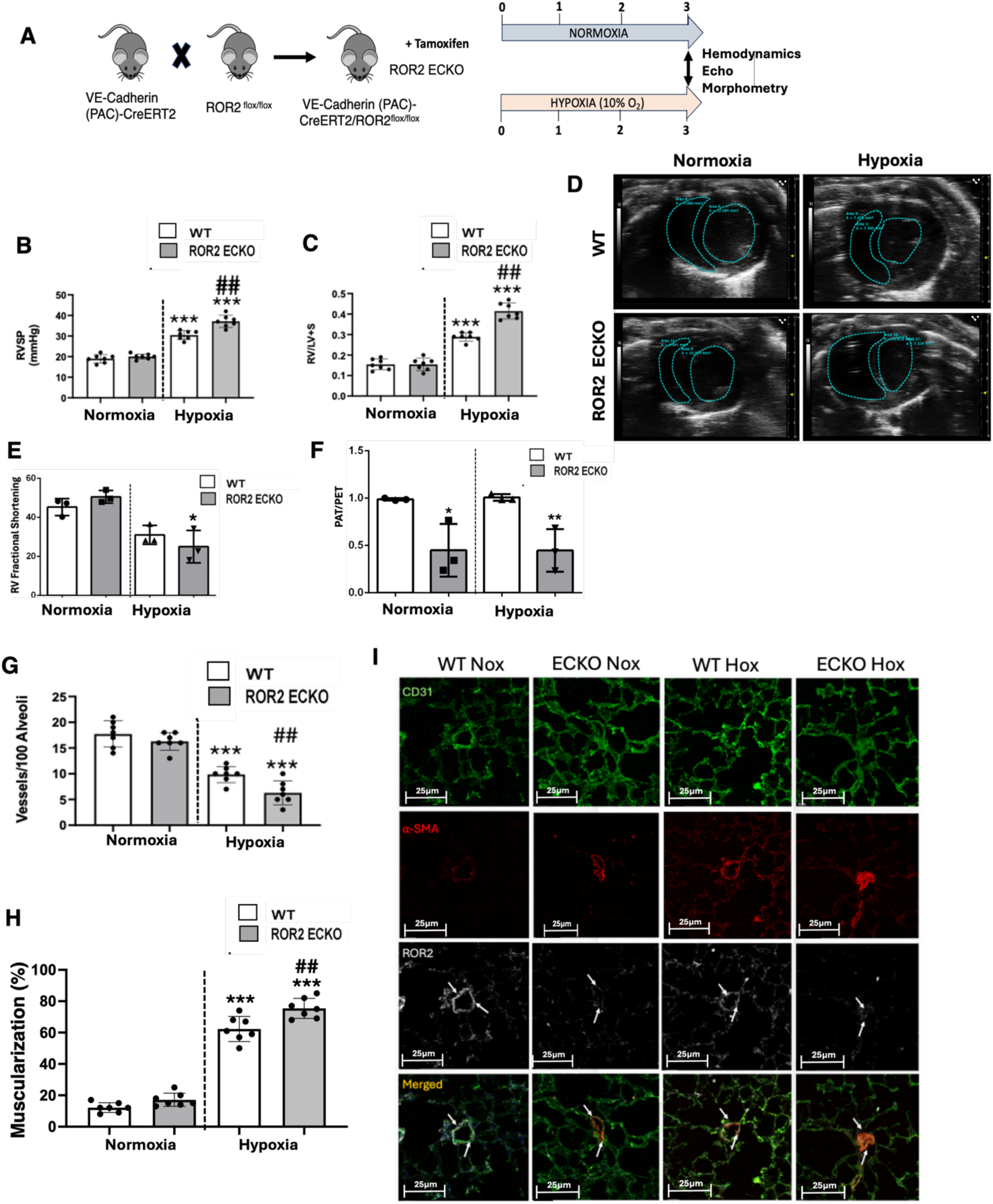
ROR2 ECKO mice develop more severe pulmonary hypertension and RV remodeling in chronic hypoxia. A) ROR2 ECKO model generation and experimental design for hypoxia studies. B–C) Right ventricular systolic pressure (RVSP, B) and Fulton index (weight ratio of the right ventricle to the sum of the left ventricle and septum, C) of WT and ROR2 ECKO mice in normoxia and hypoxia. D) Transverse echocardiographic images of WT and ROR2 ECKO mice. Dotted lines indicate RV lumen. E) RV fractional shortening, and F) Ratio of Pulmonary Artery Acceleration Time (PAT) and Pulmonary Artery Ejection Time (PET). *P<0.01, unpaired t-test. G) Vessel number per 100 alveoli and H) percent muscularization of microvessels in WT vs. ROR2 ECKO. ***: p<0.001; one-way ANOVA with Bonferroni’s post hoc test; ##P<0.01 vs corresponding WT group. I) Confocal images of endothelium (green), α-SMA (red) smooth muscle cells, and ROR2 (white) in lungs of WT and ROR2 ECKO in normoxia and chronic hypoxia.

Morphometric analysis of lung sections to assess differences in vessel number and muscularization revealed that compared to normoxia, significant reduction in vessel number was observed in both WT and ROR2 ECKO mice in hypoxia, although the severity was greater in the latter (**Fig. 1G**). Percentage of muscularized small vessel muscularization was also increased in ROR2 ECKO in normoxia and was significantly higher in hypoxia ROR2 ECKO vs. WT (**Fig. 1H**). Co-staining of αSMA (smooth muscle cells) and CD31 (Endothelial cells) in ROR2 ECKO mice shows greater medial wall thickening of the small pulmonary arteries in both normoxia and hypoxia, although severity was greater in the latter **(Fig. 1I)**. As expected, ROR2 expression was mostly absent in the endothelium of lung microvessels in ROR2 ECKO vs. WT mice (**Fig. 1I, third row**). We conclude that endothelial ROR2 insufficiency increases severity of pulmonary hypertension and vascular remodeling in response to hypoxia.

### Lung single-cell RNA sequencing (scRNA-seq) reveals a distinct gene signature in endothelial cells from ROR2 ECKO mice

To gain an understanding of cellular and molecular mechanism of ROR2 insufficiency as a contributing factor of PAH and vascular remodeling in lung endothelial cells (ECs) in mice, we performed whole lung single cell RNA-sequencing (scRNA-seq) analysis of WT and ROR2 ECKO mice under normoxia and hypoxia conditions, respectively **(Supp. Fig. 1A-G**). We further characterized the endothelial cell subpopulation as general capillary ECs (gCap), aerocytes (aCap), arterial endothelial cells 1 (AEC 1), AEC 2, and venous endothelial cells (VEC) (**Fig. 2A**). Compared to WT mice, ROR2 ECKO demonstrated a higher proportion of aCap cells in normoxia and hypoxia (**Fig. 2B**). We performed differential gene set enrichment analysis in EC subpopulation to identify candidate molecular mechanisms that could explain the phenotype of the ROR2 ECKO mice. Gene Ontology (GO) analysis for biological processes (BP) in normoxic ROR2 ECKO mice showed enrichment for genes involved in angiogenesis, and cell survival accompanied by a reduction in vesicular trafficking and immunity-related processes (**Fig. 2C**). In hypoxia, ROR2 ECKO mice demonstrated upregulation of genes associated with angiogenesis, cytoskeletal and protein homeostasis; interestingly, many of the downregulated pathways were associated with immune related processes (**Fig. 2D**). To help identify cell-related mechanisms affected by ROR2 ECKO, we performed GO cell component (CC) analysis (**Fig. 2E-F**). We found enrichment for genes associated with cytoskeleton, focal adhesions, actin filaments and contractility in ROR2 ECKO under normoxia (**Fig. 2E**). Interestingly, hypoxic ROR2 ECKO mice also demonstrated enrichment for focal adhesion and microtubule (**Fig. 2F**), which was further assessed using dot plot analysis of genes associated with cell adhesion and integrins, (**Supp. Fig. 1H-I**).

**Figure 2.**
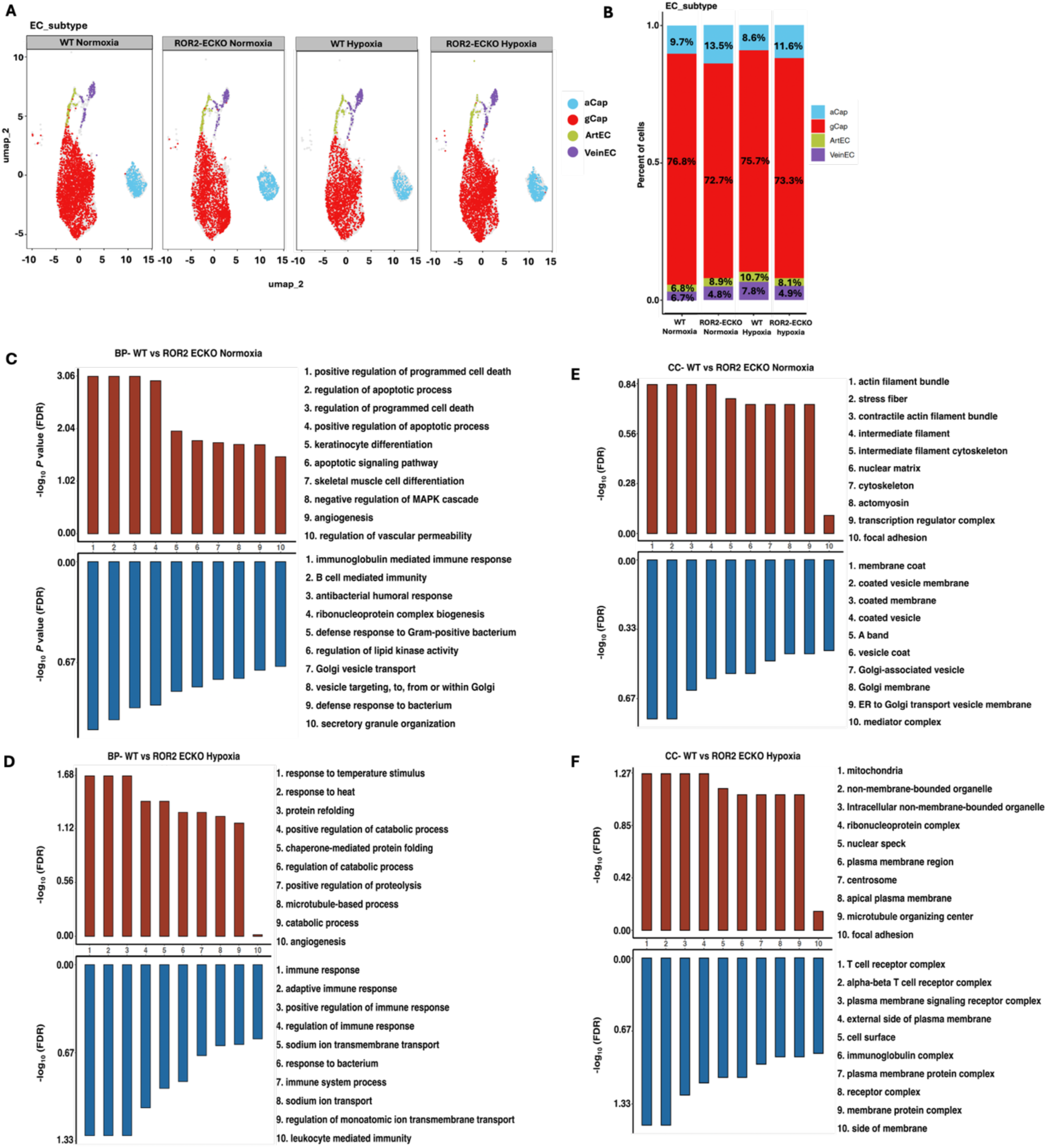
Single-cell RNA-sequencing analysis demonstrates dysregulation of genes associated with EC adhesion and barrier formation in ROR2 ECKO mice. A) Representative UMAP plot showing the distribution of the four EC subtypes (aCap, gCap, ArtEC, VeinEC) subpopulations in WT and ROR2 ECKO under normoxia and hypoxia. B) Percentage distribution of EC subtypes in WT and ROR2 ECKO lungs under normoxia and hypoxia. (C-F) GO enrichment analysis for biological process (BP, C-D) and cellular compartment (CC, E-F) of ROR2 ECKO mice vs. WT under normoxia and hypoxia.

Focal adhesions are cell membrane structures that connect the extracellular matrix to the cytoskeleton. Establishment of focal adhesions helps orchestrate the formation of the endothelial barrier, a key step in blood vessel maturation^18,19^. Previous studies shows that impairment of vascular remodeling is significantly associated with endothelial barrier dysfunction in PAH^3,20,21^. Given the findings of our scRNA-seq, we decided to look at endothelial barrier integrity in WT and ROR2 ECKO mice.

### ROR2 ECKO mice exhibit increased lung vascular leakage

To assess endothelial barrier integrity in WT and ROR2 ECKO mice, we performed Evans blue dye injection via the IVC. Under normoxic conditions, ROR2 ECKO lungs showed evident extravascular dye leakage, whereas no leakage was observed in WT lungs (**Fig. 3A**). Following hypoxic exposure, WT lungs exhibited mild dye leakage; however, ROR2 ECKO lungs demonstrated significantly greater extravasation (**Fig. 3A**). VE-cadherin, an endothelial-specific adhesion protein that maintains vascular integrity by promoting cell–cell junctions and regulating barrier function, was decreased in ROR2-ECKO vs WT mice lungs in both normoxia and hypoxia condition **(Fig. 3B).** Taken together, these studies support that ROR2 insufficiency can disrupt endothelial barrier formation, which may explain the increased severity of vascular changes seen in hypoxic ROR2 ECKO mice.

**Figure 3.**
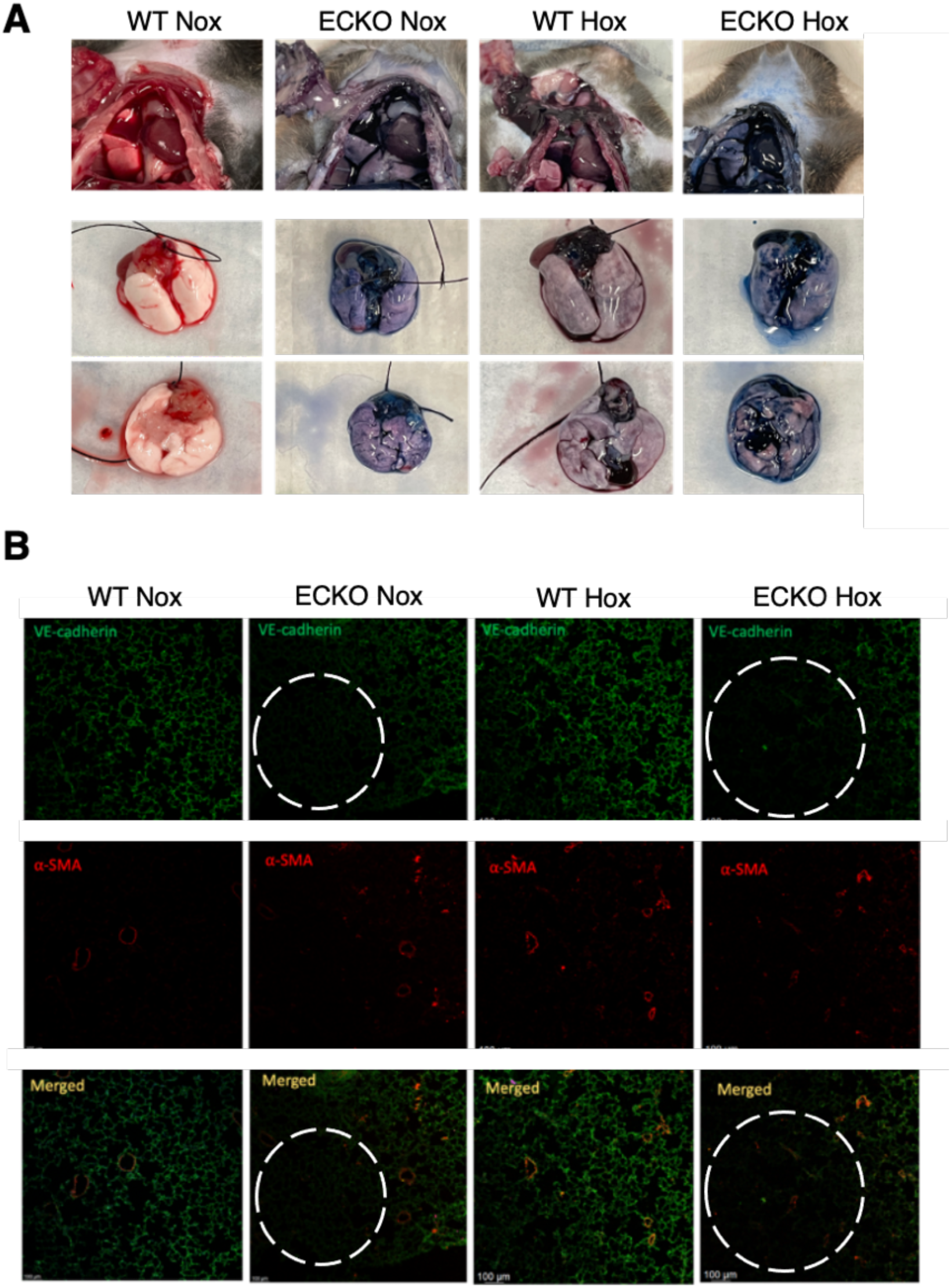
ROR2 ECKO mice demonstrate increased vascular permeability and reduced VE-cadherin expression. (A) Representative images of WT and ROR2 ECKO mice lungs after administration of Evans blue. (B) Confocal images of normoxia and hypoxia WT and ROR2 ECKO mice stained for VE-cadherin (green), and α-SMA (red). Scale bar: 100um.

### Loss of ROR2 in PMVECs is associated with increased endothelial barrier permeability and cell adhesion

To validate our *in vivo* findings in human tissue, we carried out adhesion and permeability assays in human donor PMVECs transfected with either non-targeting or ROR2-targeting (siROR2) siRNA. FITC-Dextran bead permeability assay demonstrated that, after six hours of incubation, siROR2 PMVECs demonstrated higher fluorescence rates vs. controls, indicating barrier hyperpermeability (**Fig. 4A**). Adhesion studies revealed that siROR2 PMVECs exhibited a greater number of adhered cells after one hour of seeding on a fibronectin substrate (**Fig. 4B**).

**Figure 4.**
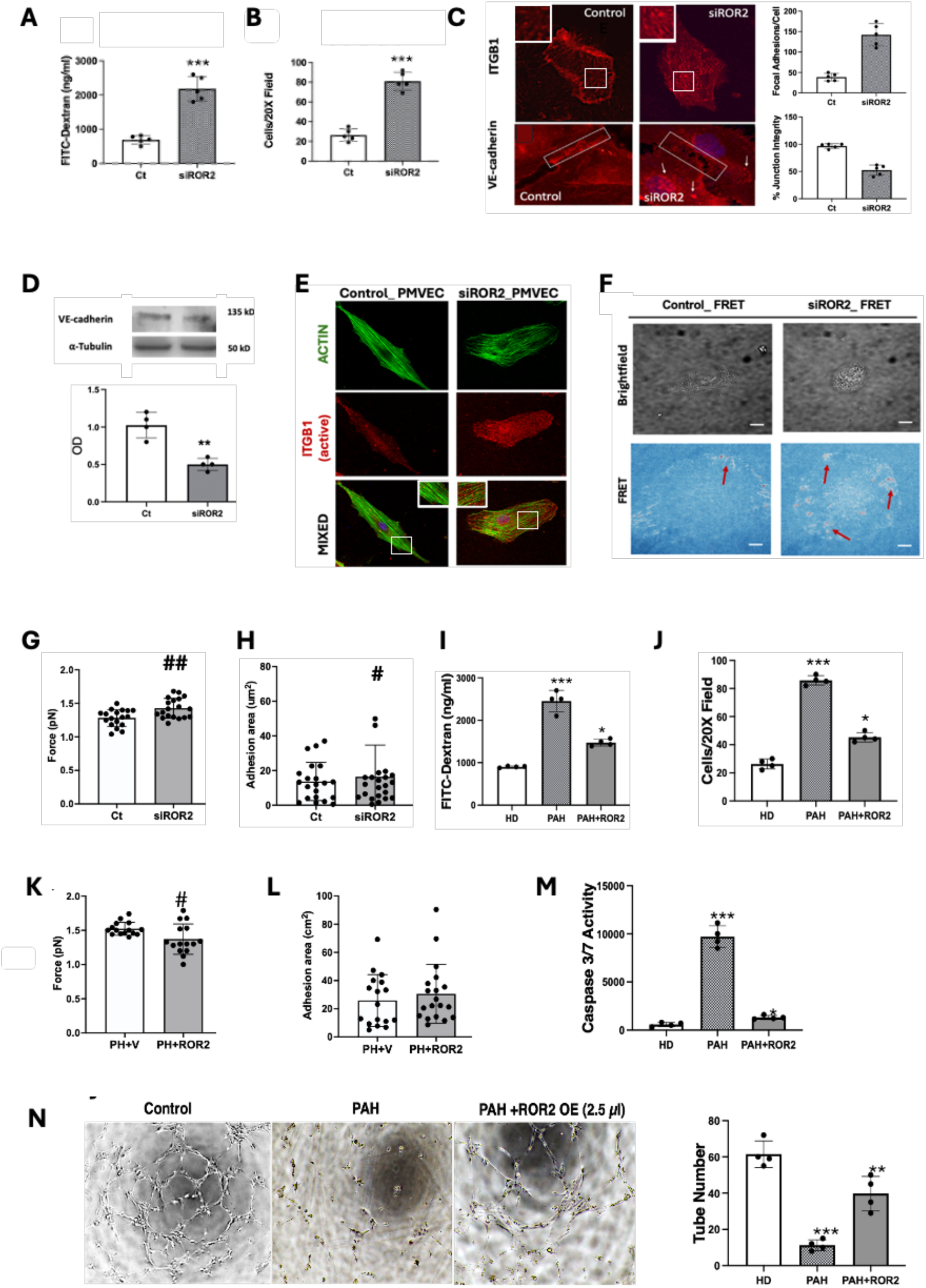
ROR2 knockdown in PMVECs increases FA density and reduces junctional integrity. A) FITC-Dextran permeability, and B) Cell adhesion assay of control and siROR2 PMVECs. ***p<0.001, unpaired t-test. C) Confocal images of ITGB1 (top) and VE-cadherin (bottom) in control and siROR2 PMVECs. White box contains a representative cell junction; arrows indicate cytoplasmic VE-cadherin granules. ***p<0.001, unpaired t-test. D) WB of VE-cadherin in control vs. siROR2 PMVECs, **p<0.01. E) Confocal imaging of control vs. siROR2 PMVECs stained for actin (green), and active ITGB1 (red). Inset contains a magnification of the area indicated by the white square. F) Representative brightfield (top) and FRET (bottom) sensor signal distribution in control and siROR2 PMVECs. Arrows indicate areas with FRET activation, indicating force generation against substrate. G-H) Average traction force per FA (G), and cell adhesion area (H) calculated from FRET signal. ##P<0.01, #P<0.05, unpaired t-test. I) FITC-Dextran and, J) Cell adhesion assays. ***P<0.001, *P<0.05, one way ANOVA with Bonferroni’s post hoc test. K) Average traction force per FA calculated from FRET signal in PAH PMVECs with ROR2 OE plasmid construct (2.5μg) compared to PAH PMVECs. #P<0.05. L) Adhesion area, P=NS. M) Caspase 3/7 apoptosis assay. ***P<0.001, *P<0.01, one way ANOVA with Bonferroni’s post hoc test. N) Representative images of Matrigel assay for healthy, PAH and PAH with ROR2 OE plasmid (2.5μg) PMVECs, respectively. Tube number was measured using ImageJ. ***P<0.001, **P<0.01, one way ANOVA with Bonferroni’s post hoc test.

Endothelial barrier formation relies heavily on the dynamic balance among focal adhesions (FAs), cytoskeletal remodeling, and intercellular communication^18,19^. This coordination integrates cellular adhesion, cytoskeletal dynamics, and cell–cell interactions, with the crosstalk between integrin complexes at FAs and VE-cadherins at adherens junctions playing a central role^22,23^. Disturbances in the balance of FA and adherens junctions could impede the establishment of the endothelial barrier, potentially precipitating vascular dysfunction and subsequent loss. Confocal microscopy demonstrated that siROR2 PMVECs shows increased number of FA clusters enriched with integrin β1 (ITGB1), a key component of the FA (**Fig. 4C, upper panel**). In contrast, adherens junction integrity was lower in siROR2 PMVECs, as evidenced by lower VE-cadherin density and gaps at cell junctions (**Fig. 4C, lower panel**). WB of whole lysates confirmed that VE-cadherin content was significantly reduced in siROR2 PMVECs vs. controls (**Fig. 4D**).

### ROR2 knockdown results in ITGB1 overactivation and cytoskeletal changes

The status of ITGB1 conformation is important in regulation of integrin activity and have a direct effect on cell adhesion, migration, and signaling. ITGB1 exists in either an inactive (i.e., extracellular domain is folded, and the ligand site is hidden) or active (extended extracellular domain, ligand site accessible). The active conformation is triggered by exposure to substrates in the extracellular matrix (i.e., fibronectin, the specific substrate of ITGB1^24^), and mechanical forces that induce changes in the cytoskeleton. ITGB1 activation also signals the recruitment of actin filaments to the cytoplasmic tail, allowing force transmission by FAs^25,26^.

To assess whether ROR2 knockdown alters ITGB1 conformational status and the actin cytoskeleton in PMVECs, we stained control and siROR2 PMVECs with an antibody specific to the active conformation of ITGB1 (clone 12G10) and phalloidin, a dye that selectively labels filamentous actin. In control PMVECs, some focal adhesions (FAs) exhibited active ITGB1 alongside well-organized actin stress fibers spanning the cell axis (**Fig. 4E, left panel**). In contrast, siROR2 PMVECs displayed widespread ITGB1 activation, with actin fibers clustered around numerous FAs, suggesting enhanced interaction between ITGB1 and the actin cytoskeleton (**Fig. 4E, right panel**).

Similar to siROR2 PMVECs, PAH PMVECs exhibit a high density of focal adhesions (FAs) across the cell membrane (**Supp. Fig. 2A**) and reduced VE-cadherin expression (**Supp. Fig. 2B**). Notably, ITGB1 within the FAs of PAH PMVECs is predominantly in an active conformation and surrounded by actin filaments, displaying a pattern reminiscent of that observed in siROR2 PMVECs. Confocal imaging further revealed that active ITGB1 is abundant in PAH vascular lesions (**Supp. Fig. 3**) and in the remodeled vessels of ROR2 ECKO mice (**Supp**. Fig. 4). It is worth noting that scRNAseq of ROR2 ECKO lungs demonstrated that, compared to WT, gCap and ArtEC exhibit the highest rates of ITGB1 expression (**Supp**. Fig. 2H).

### ROR2 deficiency increases FA traction force generation

FA activation results in increased physical communication between the cytoplasmic end of active integrins in the FA and the underlying cytoskeleton. Upon binding the ECM, the extracellular domain of ITGB1 undergoes a conformational change that regulates strength of FA binding to ECM^27,28^. To determine whether ROR2 deficiency increases ITGB1 traction force, we used a Foster Resonance Energy Transfer (FRET) ITGB1 sensor that allows visualization of dynamic ITGB1 conformational changes together with estimation of force generated at each FA **(Fig. 4F)**. Compared to controls, siROR2 PMVECs show higher traction force per FA **(Fig. 4G)** and increased adhesion area **(Fig. 4H)**. Similarly, PAH PMVECs also demonstrated higher traction force per FA, although no significant difference was found in adhesion area **(Supp. Fig. 5)**.

### Restoring ROR2 expression in PAH PMVECs improves adhesion, permeability and endothelial function

To determine whether restoring ROR2 to physiologic levels in PAH PMVECs can improve endothelial function and permeability, we transfected cells with a WT ROR2 expression construct (ROR2+) which led to an improvement in ROR2 expression compared to baseline PAH PMVECs (**Supp.** Fig. 6). Compared to vector alone, ROR2+PAH demonstrated a significant reduction in permeability (**Fig. 4I**), and adherence (**Fig. 4J**). FRET results demonstrated that ROR2 transfection reduced the force generated by FA (**Fig. 4K**) without changes in the adhesion area (**Fig. 4L**). Further evidence of the beneficial effect of ROR2 restoration was observed in survival and tube formation assays which established that, compared to vector alone, ROR2+PAH PMVECs demonstrated improved survival (**Fig. 4M**) and tube formation in Matrigel (**Fig. 4N**). Taken together, our studies confirm that ROR2 is an essential player in the regulation of barrier formation and cell adhesion and its impact translates into maintaining pro-angiogenic activities in PMVECs.

### ROR2 colocalizes with ITGB1 in FA and regulates Src-FAK phosphorylation

As the principal components of FAs, integrins are pivotal in various cellular functions, including cell migration, proliferation, differentiation, and survival. Upon achieving an active conformation, integrins become signaling hubs which can engage in two types of signaling activity. The most common is “outside-in” signaling, in which binding of extracellular matrix proteins to the ITGB1 external domain triggers downstream activation of the Src/Focal adhesion kinases (FAK) complex^29^. The Src/FAK complex comprises two intracellular tyrosine kinases that associate with integrin clusters to activate signaling pathways regulating formation of FA, adherens junctions, and cell homeostasis^30^. Following integrin activation, Src becomes phosphorylated (p416Y-Src) and recruits FAK to form a complex around integrin cytoplasmic tails. Once bound to p-Src, FAK becomes phosphorylated at tyrosine residue 397 (pY397-FAK), which results in activation and downstream signaling^31^. Studies show that loss of Src/FAK activation results in reduced survival, inadequate FA formation, and adherens junction disassembly^32,33^. Based on our observations concerning ROR2 and FA dynamics, we sought to characterize how Src/FAK signaling activity is altered by ROR2 deficiency.

As an initial step, we assessed the spatial relationship between ROR2 and ITGB1 within focal adhesion (FA) complexes. We observed strong colocalization between ROR2 and ITGB1 in control PMVECs, suggesting that ROR2 is a component of the FA complex (**Fig. 5A**). In contrast, PAH PMVECs exhibited limited colocalization, likely due to reduced ROR2 expression (**Supp. Fig. 7**).

**Figure. 5:**
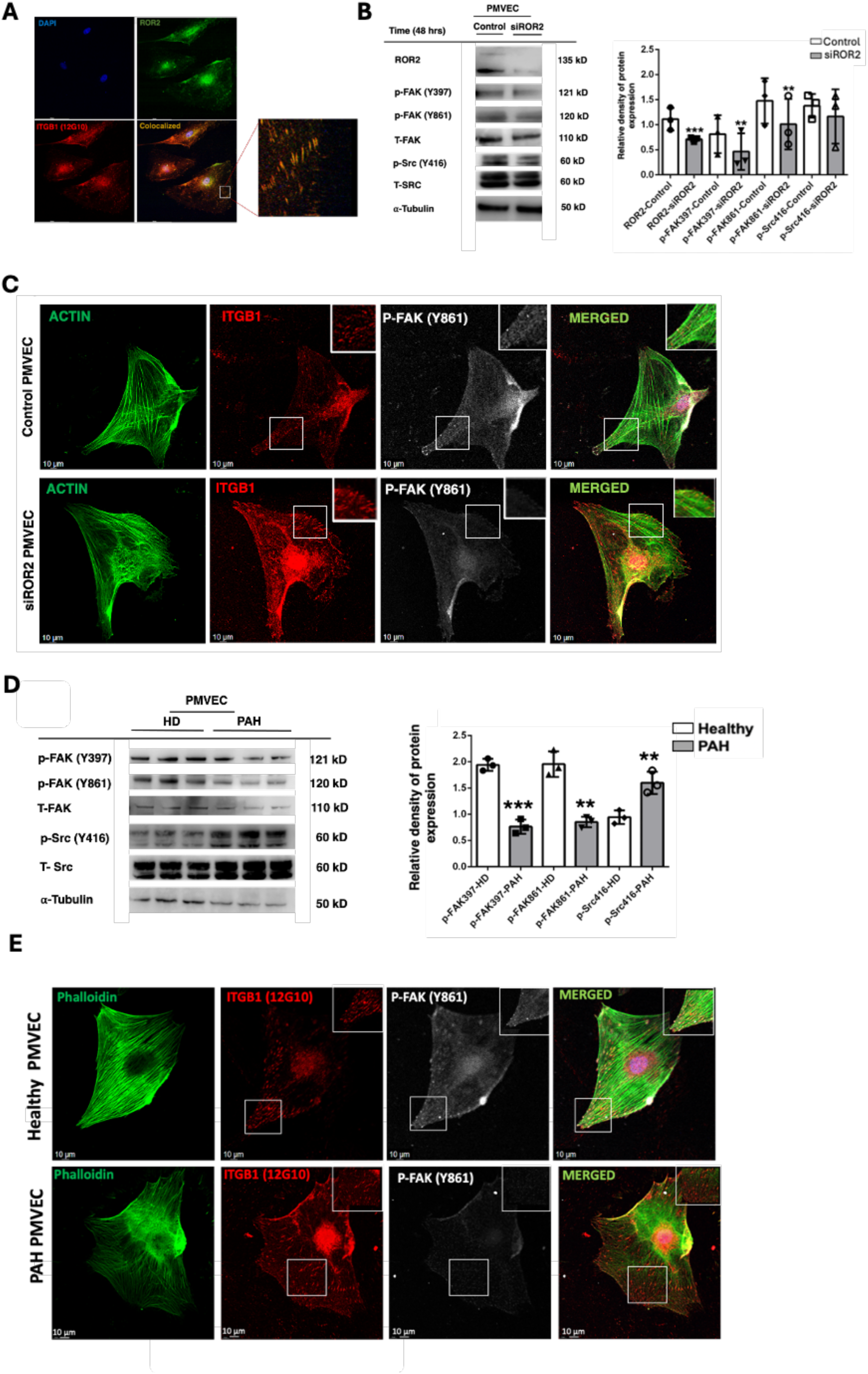
ROR2 deficiency results in uncoupling of Src/FAK phosphorylation from ITGB1 activation in PMVECs. A) Confocal imaging of PMVECs stained for ROR2 (green) and active ITGB1 (red). Magnified area demonstrates colocalization of ROR2 and active ITGB1 in FAs. B) WB of phospho (p) and total (T) FAK and Src. Densitometry relative to tubulin. ***p<0.001, **p<0.01 vs. control. C) Confocal imaging of active ITGB1 (12G10) (red), actin (green), and p-FAK (Y861, grey) for control and siROR2 PMVECs. D) WB of P/T-FAK and Src of HD and PAH PMVECs. Densitometry relative to tubulin. ***p<0.001, **p<0.01 vs. control. B) Confocal imaging of active ITGB1 (12G10) (red), actin (green), and p-FAK (Y861, grey) for HD and PAH PMVECs. Magnified area illustrates distribution of ITGB1 relative to p-FAK and actin.

Next, we examined the levels of non-phosphorylated (inactive) and phosphorylated (active) FAK (Y397/Y861) and Src (Y416) in control and siROR2 PMVECs. Despite an increased number of FAs, Western blot analysis of siROR2 PMVECs revealed significantly reduced levels of phosphorylated FAK at Y397 and Y861, indicating impaired FAK activation (**Fig. 5B**). These results were confirmed by confocal microscopy, which also showed a notable decrease in Y861-phosphorylated FAK despite elevated FA density and ITGB1 activation (**Fig. 5C**). Consistent with these findings, ROR2 ECKO mouse lungs demonstrated reduced p-FAK (Y397) levels compared to wild-type controls (**Supp. Fig. 8**).

Given that paxillin is a key downstream target of the Src/FAK signaling axis, we also evaluated phosphorylated paxillin (P-paxillin) distribution. In control PMVECs, ROR2 and P-paxillin colocalized at the cell periphery, whereas siROR2 PMVECs showed markedly reduced P-paxillin levels, further supporting diminished Src/FAK signaling (**Supp. Fig. 9**).

Finally, like siROR2 PMVECs, PAH PMVECs exhibited significantly decreased p-FAK (Y397/Y861) levels by both Western blot (**Fig. 5D**) and confocal imaging (**Fig. 5E**), reinforcing the notion of impaired FAK signaling in the PAH setting.

### RNA-seq analysis of siROR2 PMVECs identifies alterations in genes associated with integrin trafficking

To gain a better understanding of how ROR2 insufficiency triggers excess ITGB1 activation and endothelial dysfunction, we performed RNA transcriptomic profiling on control and siROR2 PMVECs. Among the genes downregulated in siROR2, we found genes relevant to vesicular trafficking (Rab12, BICD2) and regulation of FA activation (FERMT2) (**Fig. 6A**). WB confirmed significant (>50%) downregulation for BICD2, FERMT2, and Rab12 in siROR2 PMVECs (**Fig. 6B**). GO analysis further confirmed the downregulation of genes involved in regulation of vesicles (**Fig. 6C**), membrane receptor activation and signaling (**Fig. 6D**). GO analysis of biological processes (**Fig. 6E**) pointed to changes in metabolic processes as well as immune processes bearing similarities with the results from the scRNA-seq of ROR2 ECKO mice (**See Fig. 2**). Finally, KEGG analysis of siROR2 PMVECs showed increased activation of pathways associated with apoptosis, autophagy, and transcriptional regulation in cancer accompanied by downregulation of pathways involved in lipid metabolism, cytokine signaling, and toll-like receptor signaling (**Supp. Fig. 10**).

**Figure. 6:**
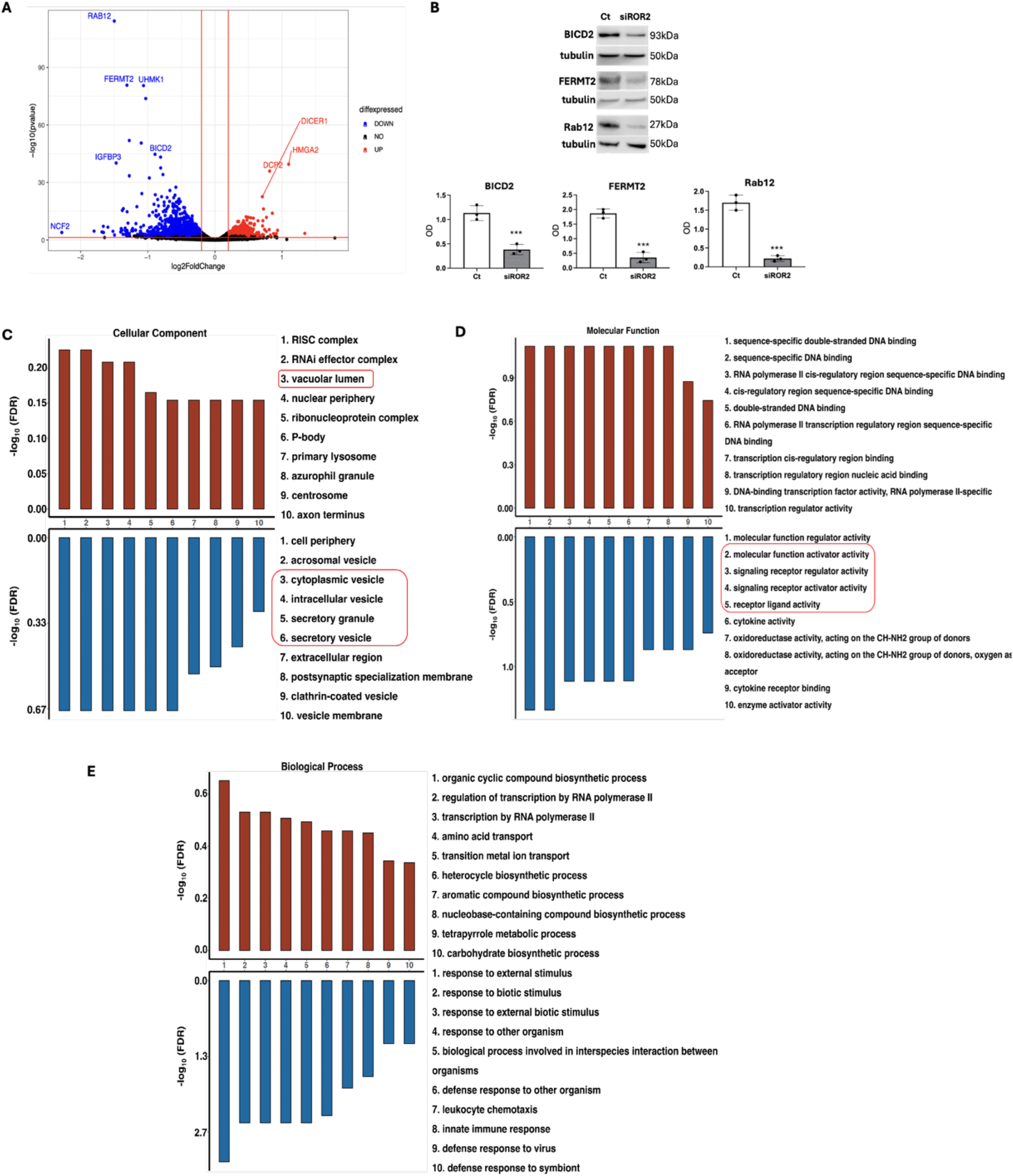
Bulk RNA-seq identifies candidate genes involved in integrin activation and vesicular trafficking in siROR2 PMVECs. A) Volcano plot map of differentially expressed genes in siROR2 vs. control PMVECs. B) WB and Densitometry of BICD2, FERMT2, and Rab12 in control and siROR2 PMVECs. ***p<0.001 vs. control; ***p<0.001 vs. HD, unpaired t-test. C-E) GO analysis for cellular compartment (C), molecular function (D), and biological process (E).

### ROR2 knockdown activates inside-out integrin signaling and alters integrin recycling

Rab12 codes for a member of the Rab GTPase family involved in integrin recycling, a process involving the endocytosis and subsequent return of integrins to the plasma membrane that is essential for regulating the availability and activity of integrins on the cell surface^34,35^. Rab12 is required to traffic vesicles towards the lysosome for proteolysis and recycling of amino acids; loss of Rab12 results in reduced vesicular trafficking thereby increasing the half-life of proteins in the membrane^36^. BICD2 codes for a critical adaptor protein that facilitates the intracellular transport of vesicles by linking them to the dynein-dynactin motor complex^37^. Another top downregulated gene was FERMT2 (i.e., Kindlin-2), which codes for kindlin-2, a cytoplasmic protein that binds to the cytoplasmic tails of β-integrin subunits, working synergistically with Talin to induce conformational changes that increase integrin affinity for ECM ligands^38^, a form of integrin signaling called “Inside-out” signaling because it does not depend on ECM binding or Src/FAK phosphorylation to trigger integrin activation.

Given the potential functional interaction of these genes in the regulation of integrin membrane abundance and activity within FAs, we hypothesized that ROR2 is involved in regulation of “Inside-out” ITGB1 activation and ITGB1 recycling. To test for “Inside-Out” integrin activity, we performed WB analysis for active ITGB1 and talin on control vs. siROR2 PMVECs. Compared to controls, siROR2 PMVECs demonstrated significantly higher active ITGB1 and Talin (∼>2-fold), supporting “Inside-Out” ITGB1 activation (**Fig. 7A**). Cell fractionation studies (**Fig. 7B**) demonstrated that, compared to controls, siROR2 PMVECs demonstrated significantly higher levels of active ITGB1 in the cytoplasm vs. membrane. In contrast, talin levels were significantly greater in siROR2 PMVEC membrane fractions. Since Rab12 is involved in integrin recycling, we compared Rab12 expression in membrane vs. cytoplasmic fractions. As expected, we found that Rab12 levels were lower in siROR2 PMVECs; however, Rab12 was higher in siROR2 cytoplasmic fractions which correlated with the higher ITGB1 observed in the siROR2 cytoplasmic extracts.

**Figure 7.**
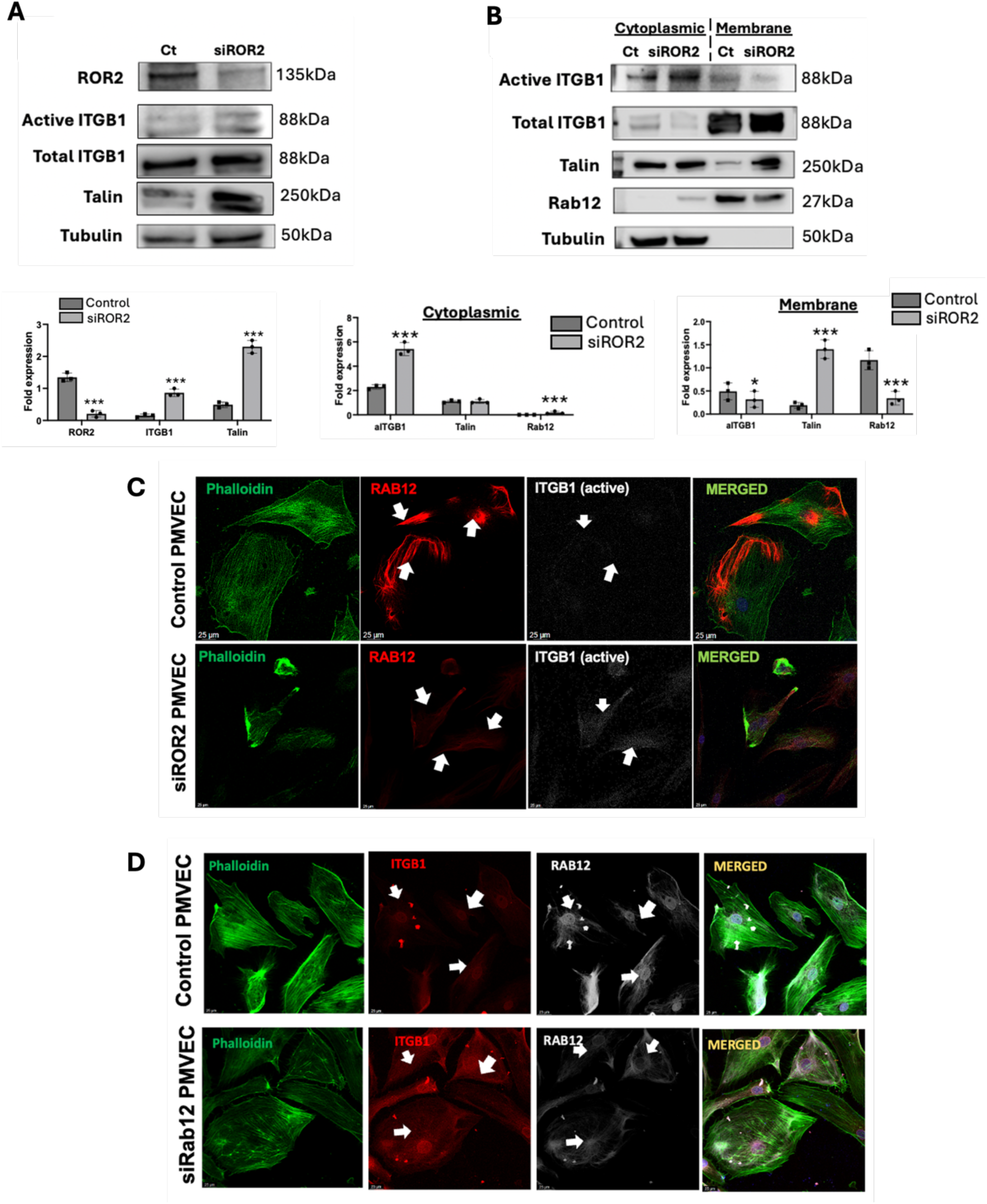
ROR2 and Rab12 regulate ITGB1 activation and recycling in PMVECs. A) WB of control (Ct) and siROR2 PMVEC whole lysates for ROR2, active and total ITGB1 and Talin. B) WB of cytoplasmic and membrane extracts from control (Ct) and siROR2 PMVECs probed against active and total ITGB1, Talin, Rab12 and tubulin. ***p<0.001 vs. control, unpaired t-test. C) Confocal imaging of control and siROR2 PMVECs probed against Rab12 (red), active ITGB1 (white) and phalloidin (actin stain, green). Arrows indicate association of ITGB1 and Rab12. D) Confocal imaging of control and siRab12 PMVECs probed against Rab12 (grey), active ITGB1 (red) and phalloidin (actin stain, green). Note contrast between low Rab12 and areas of high ITGB1 activity.

Having performed studies on siROR2 PMVECs, we sought to test whether PAH PMVECs also exhibited a similar pattern of changes in ITGB1, talin, and Rab12. In whole lysates, we found that PAH PMVECs also exhibited higher levels of active ITGB1 although the levels of Talin were only mildly increased compared to donors (**Supp. Fig. 11A**). Like the siROR2 PMVECs, cell fractionation studies showed that cytoplasmic fractions of PAH PMVECs had higher levels of active ITGB1; interestingly, we found a mild but significant increase in active ITGB1 in PAH membrane fractions. While we also found higher Talin in PAH membrane fractions, there was also significantly greater Talin protein in cytoplasmic fractions of PAH vs. donor PMVECs (**Supp. Fig. 11B).**

### Rab12 insufficiency is enough to activate ITGB1 and increase FA formation in PMVECs

Integrin recycling is tightly linked to the regulation of integrin activity, as disruption of integrin trafficking between the plasma membrane and cytoplasm can lead to inappropriate integrin activation. Noting that altered ITGB1 localization coincided with Rab12 downregulation in both siROR2 and PAH PMVECs, we sought to further investigate the role of Rab12 deficiency in ITGB1 activation.

Confocal imaging revealed that Rab12 closely associates with actin filaments, which act as intracellular tracks to guide Rab12 trafficking in cells exhibiting lower ITGB1 expression or activation (**Fig. 7C**). In control PMVECs, Rab12 was abundantly and evenly distributed throughout the cytoplasm (**Fig. 7C, top panels**). In contrast, siROR2 PMVECs displayed reduced Rab12 levels, with a marked shift in localization toward the cell membrane and a corresponding increase in ITGB1 activation (**Fig. 7C, bottom panels**).

Finally, we tested whether knocking down Rab12 would be sufficient to activate ITGB1 independent of ROR2 expression status. To do this, we use siRab12 in healthy PMVECs followed by confocal analysis of FA and active ITGB1 as previously described. As anticipated, we found that inhibition of Rab12 expression increased ITGB1 activation and FA abundance (**Fig. 7D**), without changes in ROR2 expression **(Supp.** Fig. 12).

## DISCUSSION

Endothelial dysfunction and the progressive loss of lung microvessels are significant characteristics of PAH development. Efforts have focused on how early stages of angiogenesis are altered, but few studies have examined whether the later stages of angiogenesis are also affected in PAH. The current study builds on our prior observation that Wnt7a/ROR2^11^ regulate early angiogenesis and has discovered a new role for ROR2 as a master regulator of lung endothelial adhesion and barrier formation in the late stages of angiogenesis. To the best of our knowledge, this is the first study to demonstrate a link between ROR2 and integrin signaling, spanning the formation of PMVEC focal adhesions and cell-cell communications. Based on our data, we propose a model in which ROR2 regulates the activation of ITGB1 in focal adhesions through modulation of talin-dependent “inside-out” signaling and active integrin trafficking (**Fig. 8**). Our studies are in line with the recently reported study by Lemay and colleagues demonstrating the pathological role of active ITGB1 in PAH^39^. Our study provides a mechanistic rationale for the activation of ITGB1 in PAH and supports the idea that restoring ROR2 to physiologic levels could serve as a novel therapeutic intervention to treat endothelial dysfunction and promote vascular homeostasis in the lung of PAH patients.

**Figure. 8:**
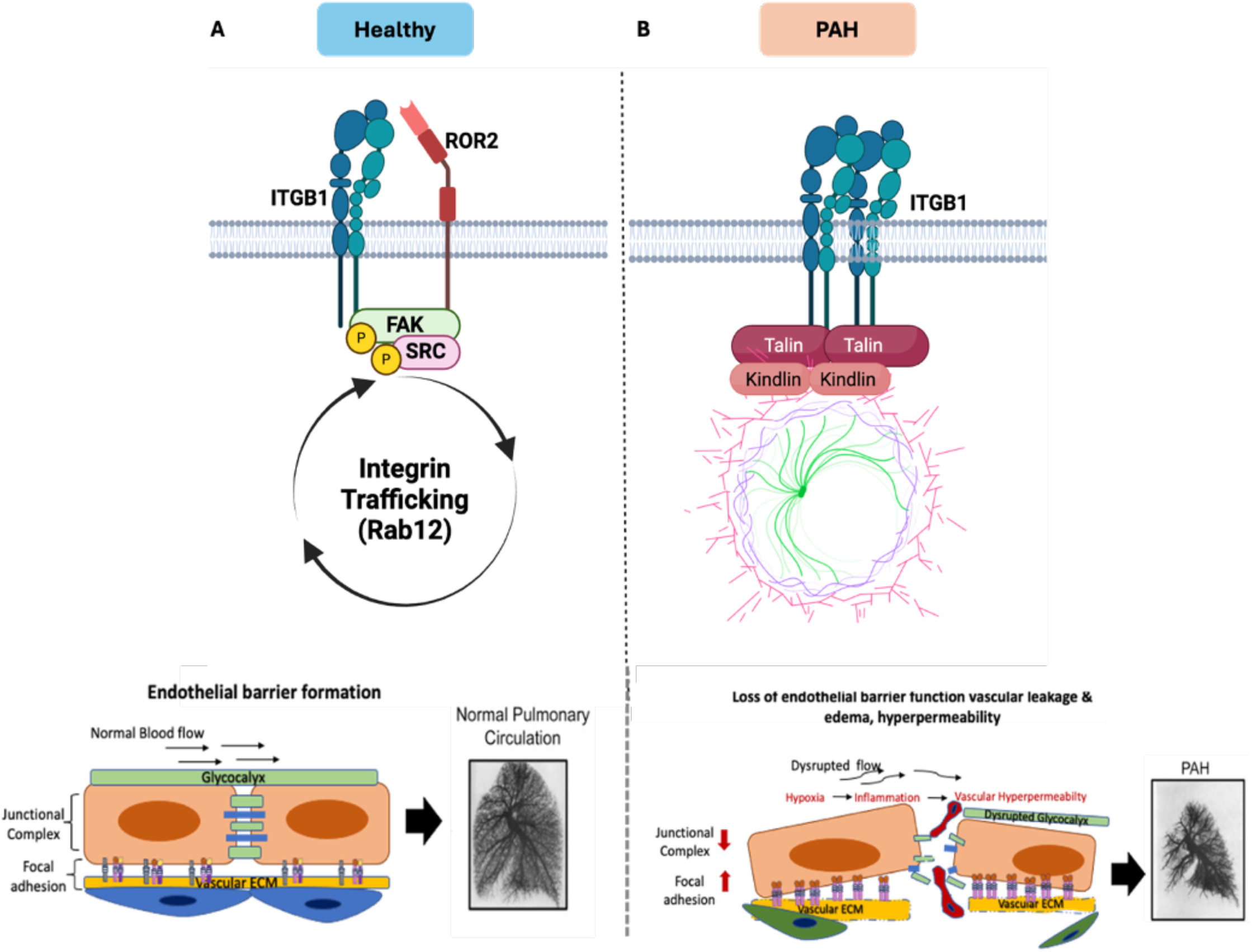
Proposed model. A) In healthy PMVECs, ROR2 interacts with ITGB1 in FA to promote Src/FAK coupling and integrin trafficking. B) Lack of ROR2 in PAH results in uncoupling of Src/FAK and Talin-dependent activation of ITGB1 resulting in excess FA and endothelial barrier dysfunction.

One of the most intriguing findings in our studies is the uncoupling of ITGB1 and Src/FAK activation. We observed that siROR2 PMVECs exhibit fewer stress fibers and actin filament disorientation, with actin clusters around ITGB1-dense areas. FAK, a non-receptor tyrosine kinase, is directly linked to the integrin cluster for activation. This integrin-associated FAK/Src pathway is crucial for cellular actin and extracellular matrix interaction and signal transmission into the cell interior^19^. The impairment in FAK/Src activation, as demonstrated in siROR2 PMVECs, results in reduced survival, inadequate FA formation, and adherens junction disassembly. The progressive loss of microvessels in the lungs is a significant feature of PAH^3^. FAK is crucial for maintaining endothelial cell survival and the integrity of the microvascular network. Impaired FAK signaling can lead to endothelial cell apoptosis and microvessel rarefaction. Previous studies in PAH and FAK activation have shed light on the complex interplay between endothelial function, vascular remodeling, and signaling pathways. FAK signaling is implicated in the proliferation and migration of pulmonary artery smooth muscle cells (PASMCs), which contribute to vascular thickening and increased resistance in PAH^40^. A recent study by Lemay and colleagues^39^ confirmed that inappropriate ITGBV/ITGB1 complex activation was a pathological feature of PAH PMVECs; however, they identified inappropriate FAK activation which was not seen in our studies. One possible explanation for the different findings is that Lemay and colleagues used endothelial cells extracted from dissected human pulmonary arteries ∼1mm in size whereas we used pulmonary microvascular endothelial cells that were extracted from whole lung using magnetic beads. Given the location and unique mechanobiology of the arteries and microvessels, it is likely that phenotypic differences exist between cell types. However, our studies agree that ITGB1 activation is a pathological event that contributes to endothelial dysfunction and should serve as the basis for treatment strategies in PAH. Our study further emphasizes that, by restoring ROR2 in PAH PMVECs, coupling of ITGB1 and Src/FAK can be restored, accompanied by the recovery of survival, adhesion, and angiogenesis. We conclude that ITGB1/FAK activity must remain within a physiologic range to support endothelial homeostasis.

Apart from integrin conformation and activation, we also observed dysregulation of vesicular trafficking and adhesion genes, as well as enriched pathways associated with vesicular trafficking, integrin-dependent signaling, cell-cell junction formation, and angiogenic pathways. Rab12 is one of the less characterized GTPases controlling intracellular trafficking between the plasma membrane and the trans-Golgi network membrane^41,42^. Previous studies have reported that Ras-associated binding guanine nucleotide triphosphatase (Rab GTPases), such as Rab5, Rab21, and Rab25, regulate integrin β1 trafficking in cancer^35^. Rab GTPases plays a multifaceted role in regulating integrin trafficking by controlling vesicle formation, intracellular sorting, endosomal maturation, membrane dynamics, and signaling crosstalk. Dysregulation of Rab-mediated integrin trafficking can lead to aberrant cell adhesion, migration, and signaling, contributing to various physiological and pathological processes, including cancer, inflammation, and vascular diseases. Rabs can crosstalk with signaling pathways involved in integrin activation and signaling, influencing integrin trafficking in response to extracellular cues. For example, Rabs may modulate the activity of kinases (e.g., FAK/Src) and phosphatases that regulate integrin function^43^. Based on our data, we hypothesize that ROR2 insufficiency in PMVECs could disrupt Rab12 activity, leading to increased ITGB1 expression and increased activation of Talin-ITGB1 which results in impaired FA formation and endothelial barrier dysfunction in PMVECs.

Our study utilized a ROR2 endothelial cell knockout (ECKO) mouse model to illustrate how ROR2 insufficiency can act as a secondary factor, exacerbating the severity of pulmonary hypertension in response to hypoxia. Interestingly, our endothelial cell (EC) knockout model of ROR2 mice exhibited significantly higher right ventricular systolic pressure (RVSP), increased RV remodeling, vessel reduction, and muscularization under chronic hypoxia. The severity of RV dysfunction in hypoxia raises questions regarding ROR2 specific effects in the cardiac endothelial function that deserve careful study in the future. We conducted permeability studies and scRNA-seq studies to further delineate how endothelial biology is disrupted in ROR2 ECKO lungs. Our data demonstrated that ROR2 insufficiency promoted endothelial dysfunction, including augmentation of angiogenesis programing, vascular permeability, impaired focal adhesion, and altered vesicle trafficking pathways in ECs which lead to disrupt vascular integrity in ROR2 ECKO mice lung. Our data are consistent with our *in vitro* ECs study. Taken together, our data indicate the importance of ROR2 in endothelial barrier formation, and lung angiogenesis, however insufficiency of ROR2 in ECs contributes to the pathogenesis of PAH development.

In conclusion, our study reveals the novel role of ROR2 in endothelial barrier formation in PMVECs, marking a pioneering discovery in this field. By modulating ITGB1 conformation and intracellular trafficking during angiogenesis in PMVECs, ROR2 deficiency profoundly impacts endothelial dysfunction, a key driver of PAH development. This novel discovery highlights the potential of selectively targeting ROR2 to restore its activity as a promising therapeutic approach in PAH.

## AUTHOR’s CONTRIBUTION

A. Mitra and V.A. de Jesus Perez was responsible for design and performance of experiments, data analysis, interpretation and drafting of the manuscript. All authors contributed to the design, performance and analysis of the studies included in the manuscript. All authors were involved in reviewing and approving the final manuscript.

## ACKNOWLEDGMENTS

Lung tissues from PAH and control patients were provided by the Pulmonary Hypertension Breakthrough Initiative, which is funded by the NIH and managed at Stanford by Drs. Marlene Rabinovitch and Roham T. Zamanian. The tissues were procured at the Transplant Procurement Centers at Stanford University, Cleveland Clinic, and Allegheny General Hospital and de-identified patient data were obtained via the Data Coordinating Center at the University of Michigan. The authors thank all patients and their proxies who participated in this study. This work utilized bioinformatics services and computing resources provided by the Stanford Genetics Bioinformatics Service Center.

## DISCLOUSER

V.A. de Jesus Perez reports support for the present manuscript from the National Institutes of Health National Heart, Lung, and Blood Institute; and outside the submitted work, holds a leadership position as AHA Chair of Diversity subcommittee. All other authors have nothing to disclose.

## Supplemental Material

Detailed methods

Supplementary tables 1-2

Supplementary Figures 1-13

Supplementary Figure legends 1-13

## NON-STANDARD ABBREVIATIONS AND ACRONYMS

PMVECs: pulmonary microvascular endothelial cells
ROR2: Receptor Tyrosine Kinase Like Orphan Receptor 2
ITGB1: Integrin beta 1
scRNA: single cell RNA sequencing
RV: right ventricle
LV: left ventricle
CreERT2: Cre recombinase (Cre) fused to a mutant estrogen ligand-binding domain (ERT2)
cKO: conditional knockout
ECKO: endothelial specific conditional knockout
DE: differential expression
BP: biological processes
CC: Cellular component
MF: Molecular Function
GO: gene ontology
GSEA: gene set enrichment analysis
NES: normalized enrichment score

